# Antimalarial drug mefloquine kills both trophozoite and cyst stages of *Entamoeba* Mefloquine and *Entamoeba histolytica*

**DOI:** 10.1101/501999

**Authors:** Conall Sauvey, Gretchen Ehrenkaufer, Anjan Debnath, Ruben Abagyan

## Abstract

*Entamoeba histolytica* is a protozoan parasite which infects approximately 50 million people worldwide, resulting in an estimated 70,000 deaths every year. Since the 1960s *E. histolytica* infection has been successfully treated with metronidazole. However, drawbacks to metronidazole therapy exist, including adverse effects, length of treatment, and the need for additional drugs to prevent transmission. All of these may decrease patient compliance and hence increase disease severity and spread of infection. In this study we identified the antimalarial drug mefloquine as possessing more potent, rapid, amoebicidal *in vitro* activity against *E. histolytica* trophozoites than metronidazole. We also showed that mefloquine could kill the cysts of a closely related reptilian parasite *Entamoeba invadens* unlike metronidazole. Additionally, mefloquine is known to possess a much longer half-life in human patients than metronidazole. This property, along with mefloquine’s rapid and broad action against *E. histolytica* position it as a promising new drug candidate against this widespread and devastating disease.

**Author Summary:** Every year, around 70,000 people worldwide die from infection by the intestinal parasite *Entamoeba histolytica*, despite the widespread availability of the drug metronidazole as a treatment. Part of the reason for this may be due to issues with patients failing to comply with the full course of treatment for the drug, due either to unpleasant side-effects, to the somewhat long treatment period, or the need for a secondary drug to kill the transmissible life stage of the parasite. In this report we discovered that the antimalarial drug mefloquine killed *E. histolytica* more potently and more rapidly than metronidazole, and, importantly, also killed the transmissible cyst stage of another *Entamoeba* species used as a model system. These findings make mefloquine an excellent candidate for an alternative drug to the current standard, with a simpler course of treatment and a more effective strategy to reduce the spread of this disease.

## Introduction

*Entamoeba histolytica* is a parasitic amoeba which infects an estimated 50 million people worldwide, resulting in around 70,000 deaths per year [1]. *E. histolytica* infection is known as amoebiasis and primarily affects the intestinal tract in humans, most commonly causing symptoms such as abdominal pain, bloody diarrhea, and colitis. In rare cases the infection spreads to other organs such as the liver and brain, and in serious cases results in patient death [2]. *E. histolytica’s* life cycle consists of a trophozoite vegetative stage which matures in its host to an infective cyst stage. The cyst stage is excreted in the host’s feces, infecting a new host when ingested via a route such as drinking contaminated water. Due to this mode of transmission *E. histolytica* disproportionately affects populations experiencing sanitation problems associated with low socioeconomic status [2–4]. Malnutrition is also known to be a major risk factor for amoebiasis, especially in children [5]. In the majority of cases where *E. histolytica* is ingested it lives asymptomatically in the human host’s intestinal tract. Symptoms develop when compromise of the mucosal layer allows it to come into contact with the intestinal wall, at which point it invades the wall and surrounding tissue causing characteristic ‘flask-shaped ulcers’ [6].

*E. histolytica* infection is currently treated with the 5-nitroimidazole drug metronidazole, which has been in use since the 1960s and experiences widespread use as a treatment against anaerobic microbial infections. However, while successful, metronidazole is not a perfect treatment against *E. histolytica*, with a few particularly notable existing issues. One of these is problems with lack of patient compliance with the course of treatment, leading to relapses and increased disease spread [7]. This is possibly due to factors such as drug adverse effects or the need for continued dosing past the resolution of disease symptoms [8]. Another issue is metronidazole’s inability to kill the infective cyst stage of *E. histolytica*. Because of this, as well as its complete absorbance from the intestines, metronidazole must be followed by a secondary luminal amoebicide paromomycin to prevent spread of the disease [9, 10]. When considered together, these factors comprise an unmet need for alternative amoebiasis therapies to metronidazole. Efforts along these lines have recently begun to be undertaken, including the development of the antirheumatic drug auranofin as a promising potential treatment for amoebiasis [11–13].

The antimalarial mefloquine is a 4-methanolquinoline compound structurally more related to quinine than chloroquine. Like metronidazole, mefloquine is a successful and widely-used antiparasitic drug. In addition to its effectiveness against *Plasmodium falciparum* and *P. vivax* mefloquine has been shown over the years to possess *in vitro* or *in vivo* activity against *Trypanosoma, Schistosoma, Echinococcosis*, and *Babesia* species [14–18]. As well as its effects on blood-stage malaria parasites, mefloquine is known for its ability to cross the blood-brain barrier, resulting in neuropsychiatric adverse events in some patients [19, 20]. Other notable pharmacokinetic attributes of mefloquine include a relatively long half-life and only partial absorption in the intestines, resulting in a profile potentially useful for persistent, invasive infections with a reservoir of parasites in the lumen [21, 22].

Based on these factors we decided to examine mefloquine for activity against *E. histolytica in vitro*. We utilized a luciferase-based cell viability assay to test potency and the rapidity of action of both mefloquine and metronidazole. We also tested mefloquine’s ability to kill the cysts of the model organism *Entamoeba invadens in vitro*. We further discuss the implications of these findings as favorable for the use of mefloquine as a clinical antiamoebic drug.

## Materials and Methods

### *E. histolytica* cell culture

*E. histolytica* strain HM-1:IMSS trophozoites were maintained in 50ml culture flasks (Greiner Bio-One) containing TYI-S-33 media, 10% heat-inactivated adult bovine serum (Sigma), 1% MEM Vitamin Solution (Gibco), supplemented with penicillin (100 U/mL) and streptomycin (100 μg/mL) (Omega Scientific) [11].

### Cell viability assay to determine drug potency against *E. histolytica*

Following a previously-published approach [11] *E. histolytica* cells, maintained in the logarithmic phase of growth were seeded into 96-well plates (Greiner Bio-One) at 5,000 cells/well to a total volume of 100 μl/well. 8- or 16-point two-fold dilution series of the treatment compounds were prepared, beginning at a maximum final treatment concentration of 50 μM. 0.5 μl of each drug concentration was added to triplicate wells for each treatment group. 0.5 μl of DMSO was used as a negative control, and 0.5 μl of 10 mM metronidazole dissolved in DMSO was used as a positive control, giving a final concentration of 50 μM. Alternatively, wells with only media were used as a negative control. The plates were placed in GasPak EZ (Becton-Dickinson) bags and incubated at 37°C for 48hr. Plates were removed and 50 μl of CellTiter-Glo (Promega) was added to each well. Plates were shaken and incubated in darkness for 20 minutes and the luminescence value of each well was read by a luminometer (EnVision, PerkinElmer). Percent inhibition was calculated by subtracting the luminescence values of each experimental data point from the average minimum signal, positive control values and dividing by the difference between the average maximum signal negative control and the positive control. The resulting decimal value was then multiplied by 100 to give a percentage.

### Trypan blue exclusion cell viability assay

*E. histolytica* trophozoites were seeded into 96-well plates at 5,000 cells/well and treated in triplicates with two-fold serially-diluted mefloquine ranging from 6.25 to 0.10 μM. Cells were incubated for 24 hr, then 10 μL of cells from each desired well were combined with 10 μl of trypan blue and the resulting mixture was counted with a hemocytometer.

### Determination of anti-amoebic drug effectiveness *in vitro* over time

Effects of mefloquine and metronidazole on *E. histolytica* trophozoite cell viability were determined as described in a previous section at a series of timepoints ranging from 0.5 to 46 hours following drug administration. EC_50_ values were calculated at each timepoint as previously described.

### Microscopic observation of drug effects on cell morphology

Confluent *E. histolytica* trophozoites were treated with 4 μM of mefloquine in 50 mL culture flasks and observed over the course of 6 hours using brightfield microscopy (Zeiss). Representative images demonstrating cellular morphology were captured at 1-hour intervals from the beginning to the end of the experiment.

### Cyst killing assay

For assays on mature cysts, a transgenic line stably expressing luciferase (CK-luc) was used [23]. Mature cyst viability assay was performed as previously descibed (Ehrenkaufer at al 2018). Parasites were induced to encyst by incubation in encystation media (47% LG) [24]. After 72 h, parasites were washed once in distilled water and incubated at 25°C for 4-5 h in water to lyse trophozoites. Purified cysts were pelleted, counted to ensure equal cyst numbers, and resuspended in encystation media at a concentration of 1-5×10^5^ cells per ml. One ml suspension per replicate was transferred to glass tubes containing encystation media and mefloquine or DMSO, then incubated at 25°C for 72 h. On the day of the assay, cysts were pelleted and treated once more with distilled water for 5 h to lyse any trophozoites that had emerged during treatment. Purified cysts were then resuspended in 75 μl Cell Lysis buffer (Promega) and sonicated for 2×10 seconds to break the cyst wall. Luciferase assay was performed using the Promega luciferase assay kit according to the manufacturer’s instructions. Assays were performed on equal volume of lysate (35 μl) and not normalized to protein content. Effect of the drug was calculated by comparison to DMSO control, after subtraction of background signal.

## Results

### Mefloquine is active against *E. histolytica* in-vitro

In order to test mefloquine for potential antiamoebic effects, an ATP-driven luciferase-based cell viability assay was employed to examine its effects on *E. histolytica* trophozoites *in vitro*. Two-fold serially-diluted concentration gradients were used to determine its 50% effective concentration (EC_50_) value. Maximum signal intensity was determined based on DMSO-treated cells, and minimum signal by cells treated with 50 μM metronidazole. From these values a cell survival percentage was calculated for each treatment dosage replicate. This experiment showed mefloquine to possess an EC_50_ value of 1.1 μM [Figure 1], notable in contrast to the previously-obtained EC_50_ value of 5 μM for metronidazole in the same *in vitro* assay system [11]. These results were replicated twice using both the same concentration range, as well as a range produced by a 1.5-fold serial dilution. Together these results suggested that mefloquine is a potent anti-amoebic agent with greater *in vitro* efficacy than the current gold-standard treatment against *E. histolytica*.

**Fig 1.**
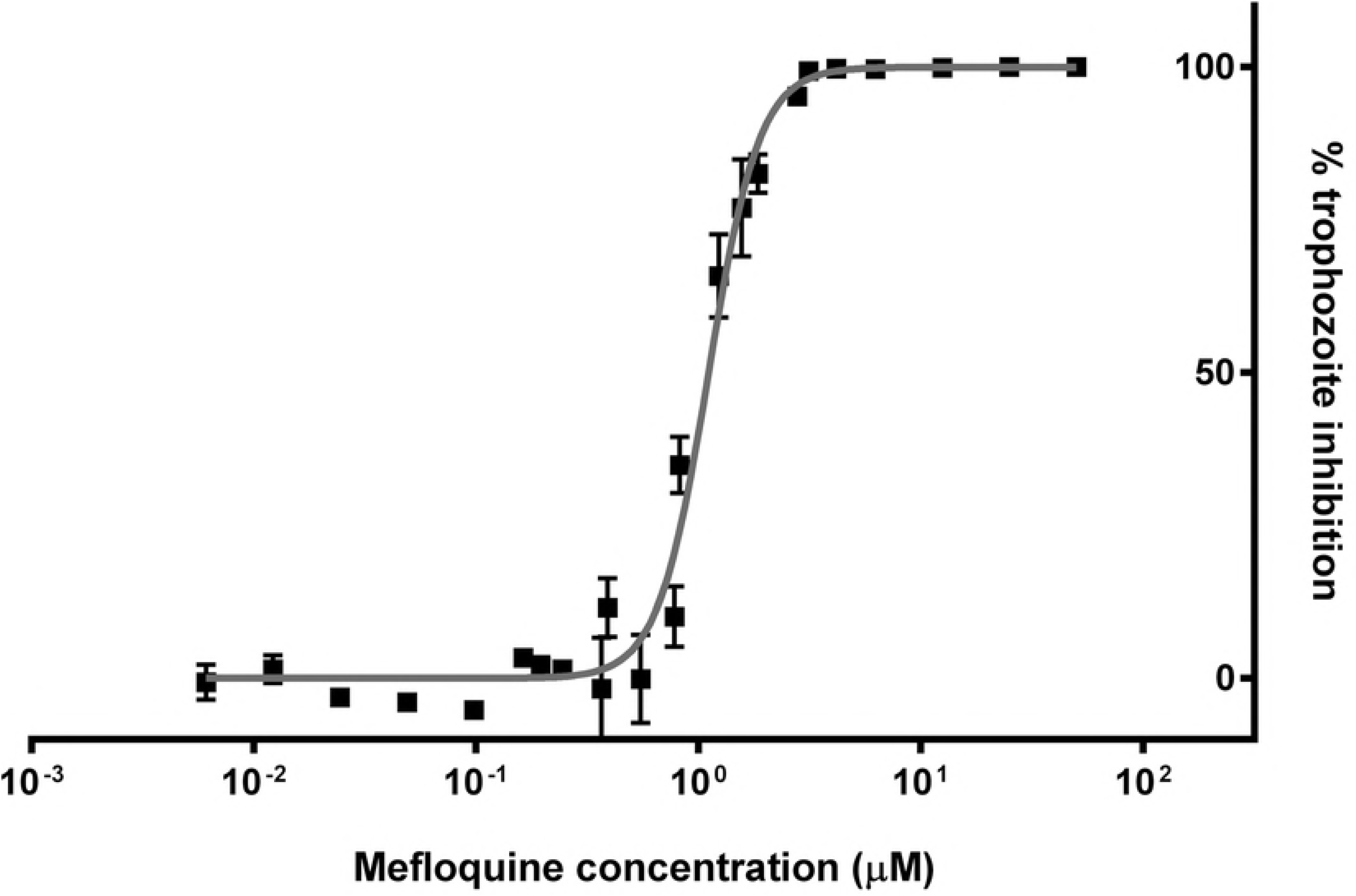
Antiamoebic activity of mefloquine. Dose response curve showing the percent inhibition of *E. histolytica* trophozoites treated with mefloquine relative to vehicle-treated control.

### Mefloquine is amoebicidal against *E. histolytica*

In order to determine whether the reduced number of viable cells in mefloquine-treated groups was due to increased cell death or merely decreased cell replication, a trypan blue exclusion assay was employed. The results of this assay confirmed the effects of mefloquine to be amoebicidal rather than amoebistatic, with the ratio of trypan-blue-permeable cells increasing with increasing drug concentration. A maximum of 100% cell staining, indicating 100% cell death, was observed at 6.25 μM of mefloquine. [Figure 2A]

**Fig 2.**
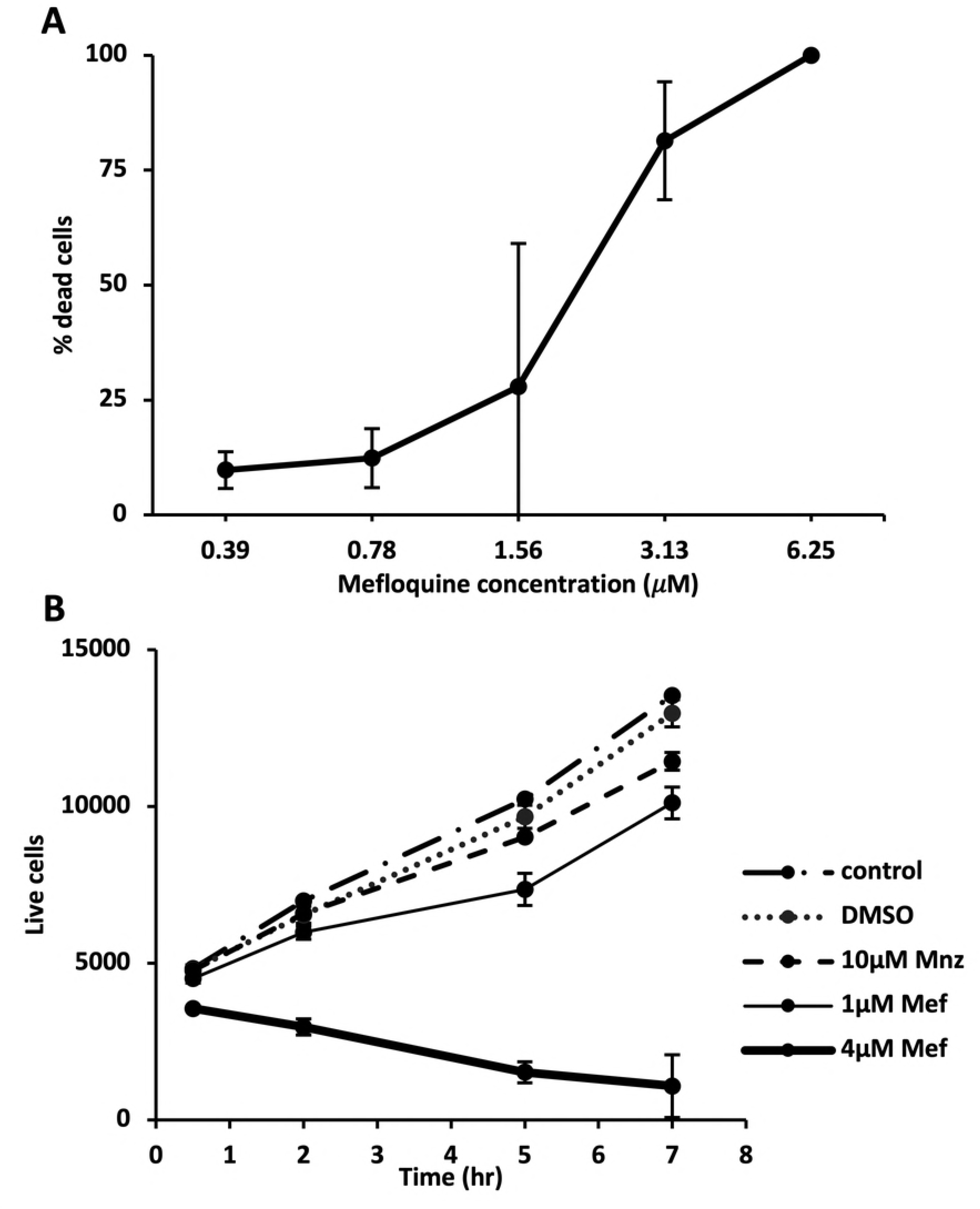
Amoebicidal activity of mefloquine. A) Percent cell death of *E. histolytica* trophozoites at 24 h in response to mefloquine treatment as quantified by trypan blue exclusion assay. B) Decline in total average trophozoite cell number per well after 4 μM mefloquine treatment as compared to either 1 μM mefloquine, 10 μM metronidazole, 0.5% DMSO, or no treatment.

The total number of live cells was observed over the course of mefloquine treatment using the same luciferase assay as previously. Replicate experimental plates were prepared and readings taken from each plate at several timepoints over the course of 7 hours.

Luminescence values obtained from control wells to which known quantities of live cells had been added were used to determine the total number of viable cells in experimental groups. In the experimental groups, trophozoites treated with 4 μM mefloquine decreased in number by a factor of five over the course of the experiments, whereas all other groups increased, including both negative controls, and those treated with 10 μM metronidazole. [Figure 2B] These results indicate that mefloquine reduces *E. histolytica* trophozoite numbers by causing cell death, rather than simply impeding or reducing replication.

### Effect of mefloquine on *E. histolytica* morphology

The morphology of *E. histolytica* trophozoites was observed after mefloquine treatment using phase contrast light microscope and compared with observations of 0.5% DMSO-treated cells. DMSO-treated trophozoites were visibly motile and irregular in shape, possessing characteristic amoeboid pseudopodia. In contrast, cells treated with 4 μM mefloquine for 6 hours were universally swollen and rounded, with dramatically enlarged vacuoles. [Figure 3]

**Fig 3.**
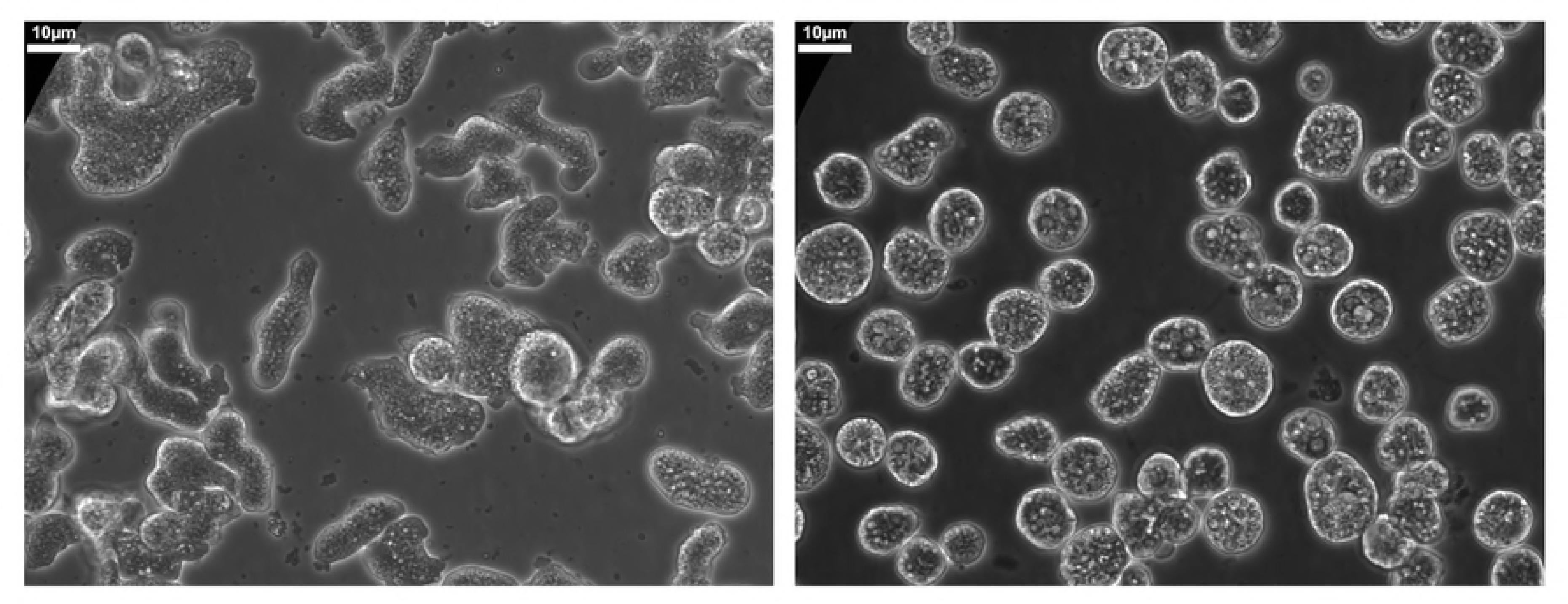
Effects of mefloquine treatment on *E. histolytica* morphology. Light microscopy images of DMSO-treated *E. histolytica* trophozoites (Left) and trophozoites treated with 4 μM of mefloquine for 6 hours (Right).

### Mefloquine kills *E. histolytica* much more rapidly than metronidazole

To further characterize the temporal aspects of the anti-amoebic effects of mefloquine, the EC_50_ values of mefloquine and metronidazole were measured at a series of timepoints after treatment of *E. histolytica* trophozoites with different concentrations of mefloquine and metronidazole. The same luciferase-based cell viability assay was used as previously to determine the percentage of cells killed in each set of experimental replicates. Duplicate plates containing cells treated with serially-diluted ranges of mefloquine concentrations were prepared for each desired timepoint. From the collected data the EC_50_ values were calculated and compared over time. Mefloquine was observed to achieve its full effectiveness as characterized by its final steady-state EC_50_ value of 1.1 μM within less than 10 hours. In contrast, metronidazole required more than 24 hours to achieve its own final value of 5 μM. [Figure 4] These results suggest that the mechanism by which mefloquine kills *E. histolytica* trophozoites is inherently more rapid than that of metronidazole.

**Fig 4.**
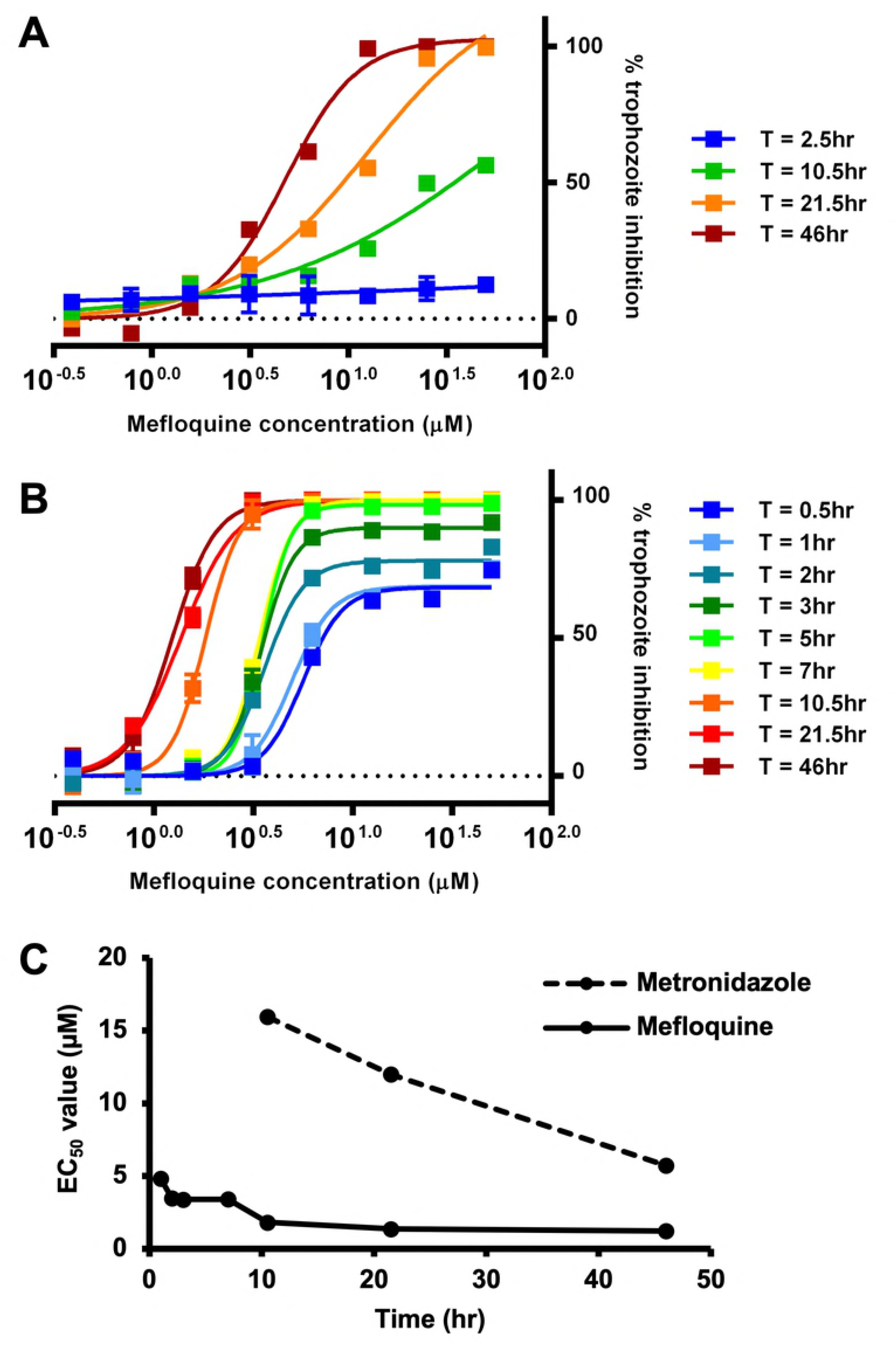
Comparison of antiamoebic activities of mefloquine and metronidazole over time. A) Dose response curves of mefloquine-treated *E. histolytica* trophozoites plotted at a series of timepoints subsequent to the start of treatment. B) Dose response curves of metronidazole-treated *E. histolytica* trophozoites plotted at a series of timepoints subsequent to the start of treatment. C) EC_50_ values for mefloquine (solid line) and metronidazole (dashed line) changing over time following the addition of each drug, eventually reaching constant values. Note: no meaningful EC_50_ value was observable in the metronidazole treatment prior to 10.5 hours

### Mefloquine kills mature *Entamoeba* cysts

A major drawback of metronidazole as a treatment for amebiasis is its poor activity against luminal parasites and cysts [2]. To determine if mefloquine may be superior in this respect, we assayed for killing of mature *Entamoeba* cysts. As *E. histolytica* cannot be induced to encyst *in vitro* [24], the related parasite, *E. invadens*, a well-characterized model system for *Entamoeba* development, was utilized. Mature (72h) cysts of a transgenic line constitutively expressing luciferase were treated with either 5 μM or 10 μM mefloquine, or 0.5% DMSO as negative control, for 3 days. After treatment, cysts were treated with distilled water for five hours to remove any remaining trophozoites, and luciferase activity was assayed. Both concentrations of mefloquine significantly reduced luciferase signal to less than 10% compared to the control (Figure 5), indicating that the drug is effectively killing the cysts. In contrast, metronidazole up to 20 μM had no effect [Figure 5].

**Figure 5.**
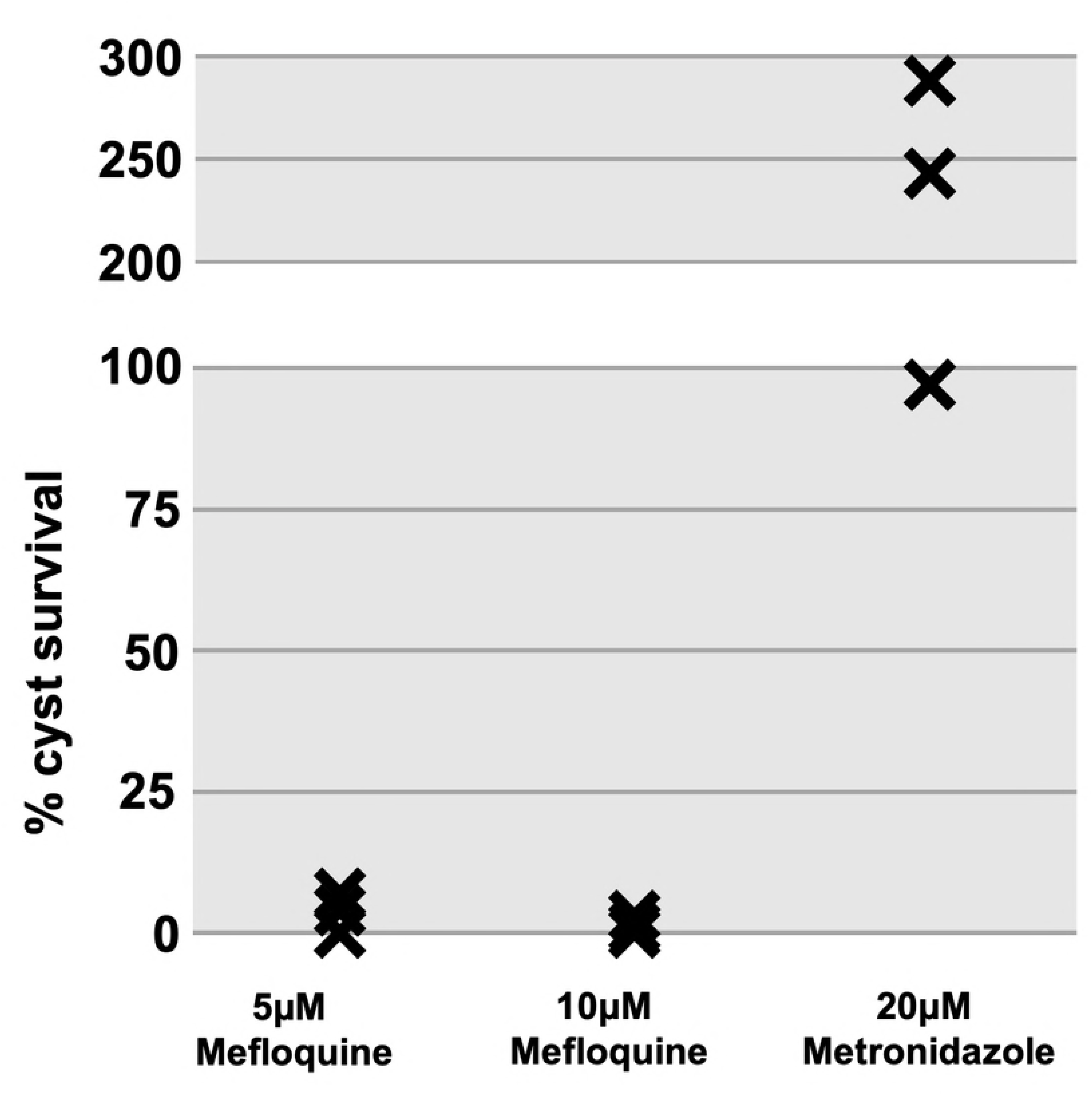
Drug activity against mature cysts. Plot displaying percentage of luciferase signal from *E. invadens* cysts treated with mefloquine or metronidazole, compared with DMSO-treated negative controls. Control readings were measured for each individual trial, which are in turn denoted by markers. metronidazole was tested at 20 μM concentration, mefloquine at both 5 μM and 10 μM.

## Discussion

In this study the FDA-approved antimalarial drug mefloquine was shown to kill *E. histolytica* trophozoites *in vitro* more potently and rapidly than the current standard therapy, metronidazole. It was also shown to kill the cyst form of related parasite *E. invadens*, which metronidazole does not. Here we will discuss how these results reveal mefloquine to have strong potential for use as an antiamoebic drug, and how in such a role it could fill gaps in the existing therapeutic paradigm for *E. histolytica* infections, reducing both the impact and spread of this disease.

The current drug of choice against *E. histolytica* infection, metronidazole, is cheap, effective, and widely used against several anaerobic pathogens [2]. However, two major concerns exist with metronidazole which render the search for additional options highly advisable.

The first concern with metronidazole therapy is patient noncompliance due to both adverse effects and the requirement for dosage past symptom improvement. Reported noncompliance has been linked with increased recurrence and prevalence of the disease and has been shown to be increased in populations known to be at greater risk for infection [8]. We found that mefloquine achieves its amoebicidal effects *in vitro* much more rapidly than metronidazole, a favorable characteristic which could result in much shorter and hence less onerous clinical dosing schedules, which in turn could partially alleviate patient noncompliance. Regarding dosage, we found a concentration of 3 μM to kill nearly 100% of *E. histolytica* trophozoites after 48 hours. Mefloquine is currently prescribed in doses of 1250 mg for cases of acute malaria, and previous studies have shown lower doses than this to be capable of producing plasma C_max_ values above the 3 μM level [21]. Additionally, the half-life of mefloquine has been shown to be up to 12 days, in contrast to a reported value of only 8 hours for metronidazole [21, 25]. All of this points to the idea that one or a small number of doses could potentially produce the desired therapeutic effects in humans that currently require many more doses of metronidazole.

The second concern is the inability of metronidazole to kill or prevent development of the infective cyst stage of *Entamoeba*. Due to its extremely high intestinal absorption, all of metronidazole’s action is systemic rather than in the lumen where a reservoir of reproducing and encysting *E. histolytica* resides. It quite effectively kills invasive trophozoites but allows the parasite’s reproductive cycle to continue, enabling the infection of other hosts. To prevent this, metronidazole therapy must be followed by treatment with a luminal antiamoebic drug such as paromomycin, which is potent in that location but not absorbed systemically at all [9, 10]. The current therapeutic strategy thus relies on the sequential administration of two separate drugs each with opposite absorption profiles in order to effectively control both the symptoms and spread of amoebiasis. Such a lengthy and complex treatment no doubt greatly aggravates the issues with patient noncompliance described in the first concern. In this study we documented features of mefloquine which might allow it to circumvent this dual-drug issue. We tested mefloquine for effectiveness against the cyst form of *E. invadens* and found it to be active. *E. invadens* is a similar parasite to *E. histolytica* and is widely used as an *in vitro* model of encystation for *E. histolytica* [26]. In this study it was found that mefloquine was capable of killing 100% of *E. invadens* cysts at 5 μM, a value close to that observed to kill *E. histolytica* trophozoites. Importantly, unlike metronidazole, around 20-30% of mefloquine remains in the intestines of patients after absorption, raising the possibility that it could act as a luminal antiamoebic against both cysts and trophozoites [22]. As such mefloquine has the potential to fill the cysticidal and amoebicidal roles required for successful treatment of *E. histolytica* infection.

Mefloquine’s systemic distribution includes the ability to cross the blood-brain barrier - a feature which has both negative and positive consequences [27]. Negatively, mefloquine is known for producing neuropsychiatric adverse effects in a subset of patients [28]. Positively, mefloquine’s ability to enter the brain allows it to act against parasites that have invaded that region. While most literature on the subject involves combinations of mefloquine with other drugs such as artesunate for treatment of clinical or experimental cerebral malaria, cases have been reported where mefloquine alone has been successful in patients [29, 30]. Like malaria parasites, *E. histolytica* invades the brain in a small number of extreme cases [31]. Mefloquine’s ability to cross the blood-brain barrier might allow it to act as an effective treatment in such cases. This would add to the overall versatility and usefulness of mefloquine as an antiamoebic drug.

Given the results of this study a key question remaining is the nature of mefloquine’s mechanism of action against *E. histolytica*. In *Plasmodium* species as well as various other cell types a number of hypotheses have been proposed. These hypotheses range from the inhibition of plasma membrane dynamics to the induction of reactive oxygen species stress [32–36]. One particular hypothesis raised by two papers from the past decade proposed that mefloquine might act against *Plasmodium* parasites through inhibition of the cytosolic ribosome and resultant suppression of protein translation [37, 38]. While these studies do support the idea that mefloquine may act at least partially via such a mechanism in *Plasmodium*, it is unclear whether ribosomal inhibition might also be lethal in *E. histolytica*. Some clues regarding the answer to this question come from existing compounds known to kill *E. histolytica*. The drug paromomycin, currently prescribed as a luminal amoebicide, is known to act through ribosomal inhibition against both bacteria and *Leishmania* species [39, 40]. Additionally, potent activity of the ribosomal inhibitor anisomycin against *E. histolytica* trophozoites and *E. invadens* cysts has recently been reported [13]. Together these drug activities confirm that compounds targeting the ribosomal complex are capable of killing *E. histolytica*, rendering it conceivable that such a mechanism might be at work with the effects of mefloquine. Future studies should explore this possibility, as well as investigate ribosome-targeting compounds as a possible class of amoebicides.

In conclusion we demonstrated *in vitro* that the FDA-approved antimalarial drug mefloquine has the potential to act as a new treatment option for *E. histolytica* infection. Mefloquine is superior to the current practice, due to greater potency, rapidity of action, and cysticidal effects. Further studies using *in vivo* models of the disease should be undertaken to refine the optimal dosage and duration of treatment.

## Acknowledgments

The authors of this study would like to thank Dr. Jim McKerrow and the Center for Discovery and Innovation in Parasitic Diseases at the Skaggs School of Pharmacy at the University of California - San Diego as well as Da Shi, Lily Hahn, and Abdolhakim Mohammed for their contributions to this work.

---

S1 Table. Figure 1 data

S2 Table. Figure 2A data

S3 Table. Figure 2B data

S4 Table. Figure 4 data

S5 Table. Figure 5 data

## References

1. Shirley DT, Farr L, Watanabe K, Moonah S. A Review of the Global Burden, New Diagnostics, and Current Therapeutics for Amebiasis. Open forum infectious diseases. 2018; 5(7):ofy161.

2. Pritt BS, Clark CG. Amebiasis. Mayo Clin Proc. 2008; 83(10):1154-9; quiz 9-60.

3. Faria CP, Zanini GM, Dias GS, da Silva S, de Freitas MB, Almendra R, et al. Geospatial distribution of intestinal parasitic infections in Rio de Janeiro (Brazil) and its association with social determinants. PLoS Negl Trop Dis. 2017; 11(3):e0005445.

4. Sahimin N, Lim YA, Ariffin F, Behnke JM, Lewis JW, Mohd Zain SN. Migrant Workers in Malaysia: Current Implications of Sociodemographic and Environmental Characteristics in the Transmission of Intestinal Parasitic Infections. PLoS Negl Trop Dis. 2016; 10(11):e0005110.

5. Verkerke HP, Petri WA, Jr., Marie CS. The dynamic interdependence of amebiasis, innate immunity, and undernutrition. Semin Immunopathol. 2012; 34(6):771-85.

6. Ralston KS, Petri WA, Jr. Tissue destruction and invasion by Entamoeba histolytica. Trends in parasitology. 2011; 27(6):254-63.

7. Dusengeyezu E, Kadima J. How do Metronidazole Drawbacks Impact Patient Compliance and Therapeutic Outcomes in Treating Amoebiasis in Rwanda. International Journal of TROPICAL DISEASE & Health. 2016; 17(3):1-7.

8. Garduno-Espinosa J, Martinez-Garcia MC, Fajardo-Gutierrez A, Ortega-Alvarez M, Alvarez-Espinosa A, Vega-Perez V, et al. Frequency and risk factors associated with metronidazole therapeutic noncompliance. Revista de investigacion clinica; organo del Hospital de Enfermedades de la Nutricion. 1992; 44(2):235-40.

9. Kikuchi T, Koga M, Shimizu S, Miura T, Maruyama H, Kimura M. Efficacy and safety of paromomycin for treating amebiasis in Japan. Parasitology international. 2013; 62(6):497-501.

10. Blessmann J, Tannich E. Treatment of asymptomatic intestinal Entamoeba histolytica infection. The New England journal of medicine. 2002; 347(17):1384.

11. Debnath A, Parsonage D, Andrade RM, He C, Cobo ER, Hirata K, et al. A high- throughput drug screen for Entamoeba histolytica identifies a new lead and target. Nat Med. 2012; 18(6):956-60.

12. Bashyal B, Li L, Bains T, Debnath A, LaBarbera DV. Larrea tridentata: A novel source for anti-parasitic agents active against Entamoeba histolytica, Giardia lamblia and Naegleria fowleri. PLoS Negl Trop Dis. 2017; 11(8):e0005832.

13. Ehrenkaufer GM, Suresh S, Solow-Cordero D, Singh U. High-Throughput Screening of Entamoeba Identifies Compounds Which Target Both Life Cycle Stages and Which Are Effective Against Metronidazole Resistant Parasites. Front Cell Infect Microbiol. 2018; 8:276.

14. Crouch AA, Seow WK, Thong YH. Effect of twenty-three chemotherapeutic agents on the adherence and growth of Giardia lamblia in vitro. Transactions of the Royal Society of Tropical Medicine and Hygiene. 1986; 80(6):893-6.

15. Planer JD, Hulverson MA, Arif JA, Ranade RM, Don R, Buckner FS. Synergy testing of FDA-approved drugs identifies potent drug combinations against Trypanosoma cruzi. PLoS Negl Trop Dis. 2014; 8(7):e2977.

16. Xiao SH. Mefloquine, a new type of compound against schistosomes and other helminthes in experimental studies. Parasitol Res. 2013; 112(11):3723-40.

17. Liu C, Zhang H, Yin J, Hu W. In vivo and in vitro efficacies of mebendazole, mefloquine and nitazoxanide against cyst echinococcosis. Parasitol Res. 2015; 114(6):2213-22.

18. Munkhjargal T, AbouLaila M, Terkawi MA, Sivakumar T, Ichikawa M, Davaasuren B, et al. Inhibitory effects of pepstatin A and mefloquine on the growth of Babesia parasites. Am J Trop Med Hyg. 2012; 87(4):681-8.

19. Greenwood BM, Fidock DA, Kyle DE, Kappe SH, Alonso PL, Collins FH, et al. Malaria: progress, perils, and prospects for eradication. The Journal of clinical investigation. 2008; 118(4):1266-76.

20. Schlagenhauf P. Mefloquine for malaria chemoprophylaxis 1992-1998: a review. Journal of travel medicine. 1999; 6(2):122-33.

21. Karbwang J, Na Bangchang K, Thanavibul A, Back DJ, Bunnag D, Harinasuta T. Pharmacokinetics of mefloquine alone or in combination with artesunate. Bulletin of the World Health Organization. 1994; 72(1):83-7.

22. Looareesuwan S, White NJ, Warrell DA, Forgo I, Dubach UG, Ranalder UB, et al. Studies of mefloquine bioavailability and kinetics using a stable isotope technique: a comparison of Thai patients with falciparum malaria and healthy Caucasian volunteers. British journal of clinical pharmacology. 1987; 24(1):37-42.

23. Ehrenkaufer GM, Singh U. Transient and stable transfection in the protozoan parasite Entamoeba invadens. Mol Biochem Parasitol. 2012; 184(1):59-62.

24. Sanchez L, Enea V, Eichinger D. Identification of a developmentally regulated transcript expressed during encystation of Entamoeba invadens. Mol Biochem Parasitol. 1994; 67(1):125-35.

25. Welling PG, Monro AM. The pharmacokinetics of metronidazole and tinidazole in man. Arzneimittel-Forschung. 1972; 22(12):2128-32.

26. Ehrenkaufer GM, Weedall GD, Williams D, Lorenzi HA, Caler E, Hall N, et al. The genome and transcriptome of the enteric parasite Entamoeba invadens, a model for encystation. Genome biology. 2013; 14(7):R77.

27. Pham YT, Nosten F, Farinotti R, White NJ, Gimenez F. Cerebral uptake of mefloquine enantiomers in fatal cerebral malaria. International journal of clinical pharmacology and therapeutics. 1999; 37(1):58-61.

28. Lee SJ, Ter Kuile FO, Price RN, Luxemburger C, Nosten F. Adverse effects of mefloquine for the treatment of uncomplicated malaria in Thailand: A pooled analysis of 19, 850 individual patients. PloS one. 2017; 12(2):e0168780.

29. Di Perri G, Olliaro P, Ward S, Allegranzi B, Bonora S, Concia E. Rapid absorption and clinical effectiveness of intragastric mefloquine in the treatment of cerebral malaria in African children. The Journal of antimicrobial chemotherapy. 1999; 44(4):573-6.

30. Sun HY, Fang CT, Wang JT, Kuo PH, Chen YC, Chang SC. Successful treatment of imported cerebral malaria with artesunate-mefloquine combination therapy. Journal of the Formosan Medical Association = Taiwan yi zhi. 2006; 105(1):86-9.

31. Petri WA, Haque R. Entamoeba histolytica brain abscess. Handbook of clinical neurology. 2013; 114:147-52.

32. Fitch CD. Ferriprotoporphyrin IX, phospholipids, and the antimalarial actions of quinoline drugs. Life sciences. 2004; 74(16):1957-72.

33. Gunjan S, Singh SK, Sharma T, Dwivedi H, Chauhan BS, Imran Siddiqi M, et al. Mefloquine induces ROS mediated programmed cell death in malaria parasite: Plasmodium. Apoptosis: an international journal on programmed cell death. 2016; 21(9):955-64.

34. Paivandy A, Calounova G, Zarnegar B, Ohrvik H, Melo FR, Pejler G. Mefloquine, an anti-malaria agent, causes reactive oxygen species-dependent cell death in mast cells via a secretory granule-mediated pathway. Pharmacology research & perspectives. 2014; 2(6):e00066.

35. Yadav N, Dwivedi A, Mujtaba SF, Verma A, Chaturvedi R, Ray RS, et al. Photosensitized mefloquine induces ROS-mediated DNA damage and apoptosis in keratinocytes under ambient UVB and sunlight exposure. Cell biology and toxicology. 2014; 30(5):253-68.

36. Yan KH, Yao CJ, Hsiao CH, Lin KH, Lin YW, Wen YC, et al. Mefloquine exerts anticancer activity in prostate cancer cells via ROS-mediated modulation of Akt, ERK, JNK and AMPK signaling. Oncology letters. 2013; 5(5):1541-5.

37. Gamo FJ, Sanz LM, Vidal J, de Cozar C, Alvarez E, Lavandera JL, et al. Thousands of chemical starting points for antimalarial lead identification. Nature. 2010; 465(7296):305-10.

38. Wong W, Bai XC, Sleebs BE, Triglia T, Brown A, Thompson JK, et al. Mefloquine targets the Plasmodium falciparum 80S ribosome to inhibit protein synthesis. Nat Microbiol. 2017; 2:17031.

39. Shalev-Benami M, Zhang Y, Rozenberg H, Nobe Y, Taoka M, Matzov D, et al. Atomic resolution snapshot of Leishmania ribosome inhibition by the aminoglycoside paromomycin. Nature communications. 2017; 8(1):1589.

40. Tok JB, Bi L. Aminoglycoside and its derivatives as ligands to target the ribosome. Current topics in medicinal chemistry. 2003; 3(9):1001-19.

